# Identification of bacteria strains using the Recombinase Polymerase Amplification assay on a miniaturized solid-state pH sensor

**DOI:** 10.1101/2022.01.04.474950

**Authors:** Anh H. Nguyen, Samir Malhotra, Michael P.H. Lau, Hung Cao

## Abstract

Rapid identification of bacteria based on nucleic acid amplification allows dealing with the detection of pathogens in clinical, food, and environmental samples. Amplification product must be detected and analyzed by external devices or integrated complicated optical systems. Here, we developed a solid-state pH electrode based on iridium oxide (IrO_2_) films to measure released hydrogen ions (H^+^) from isothermal nucleic acid (NA) amplification of bacterial samples. By recombinase polymerase amplification (RPA), we achieved rapid (< 15 min) and sensitive (<30 copies) detection with an accuracy of about 0.03 pH. The RPA-based hydrogen ion sensing assay shows higher specificity, sensitivity, and efficiency as the same polymerase chain reaction (PCR) methods. We initially used the RPA-based sensor to detect *E. coli* species in laboratory samples. Among, 27 random laboratory samples of *E. coli* samples, 6 were found to be DH5alpha, 9 BL21, 3 HB101, 6 TOP10, and 3 JM109. The electrical detection of amplification provides generally applicable techniques for the detection of nucleic acid amplification, enabling molecular diagnostic tests in the field and integrating data transmission to the mobile device. These results can be future developed into an efficient tool for rapid on-site detection of bacterial pathogens in clinical samples.

## Introduction

The prevalence of pathogenic bacterial contamination and its effect on human health has called for alternative methods to laboratory-based techniques of bacterial detection. Biosensors based on isothermal DNA amplification techniques will help solve drawbacks in on-site detections of pathogens [1]. Besides using fluorescent indicators, researchers have focused more on the use of released hydrogen ions (H^+^) from DNA amplification to detect pathogens [2-5]. The sensing H^+^ on ion-sensitive field-effect transistors (ISFET) has become the standard indicator for rapid detection [6], quantification [2], and DNA sequencing [7], which means that H^+^ from NA amplification is a reliable indicator to detect pathogens. While the metal oxide pH sensors have been used to detect pH of blood [8], skin [9], and urine [10], the sensors have not been explored as a reliable platform to detect H^+^ from NA amplification. Recently, a label-free and electrochemical detection of NA based on primer-generation rolling circle amplification were developed on the iridium oxide pH sensor [11].

Yamanaka’s protocol [12] has been used for direct anodic deposition of IrO_2_ film on gold electrode. The modified IrO_2_ electrode showed super Nernstian response from 65 ± 0.5 -72.9 ± 0.9 mV/pH with resolution of 0.03 pH units, as same as ion-sensitive field-effect transistor (ISFET), under the various pH buffer solutions. Its potential response is less effected in the presence of proteins in samples [6, 13, 14]. Moreover, mechanistic and response studies including interferences of ions of the pH sensors against sensing surface have also been investigated [15, 16]. Various surface morphologies of the electrode have been developed through electrodeposition and sputter deposition processes, the most widespread techniques for thin metallic film deposition, on different substrates [14, 17, 18]. Electrodeposition is a more cost-effective and faster technique which coat larger areas compared to sputtering and chemical vapor deposition techniques [19]. Here we electrodeposited IrO_2_ layer on planar gold electrodes to form IrO_2_-based pH sensor to detect released H+ from isothermal DNA amplification of E. coli strains. Recombinase polymerase amplification (RPA), a versatile isothermal DNA amplification, has been used in clinical diagnostics [20]. For example, RPA has been used in analytical and clinical assessment of a portable assay for urogenital *Schistosomiasis* diagnostics [21], Potato virus Y, O, and N types in potato [22] [23]. *Cryptococcus neoformans/C. gattii* in cerebral spinal fluid [24], and *Mycoplasma bovis*, and *Campylobacter jejuni* in food samples [25]. Compared to other isothermal amplification methods, RPA allows NA amplification at power saving modes (such as at room temperature), simple primer design, extremely quick (20 min), no initial heating step, and affordable to various biological samples [26].

This first-time approach was to investigate the use of solid-state pH sensor for measuring released hydrogen ions from RPA and evaluating ion interferences presenting in PCR buffers to the pH electrodes. The simple design of our chip contains a closed pair of working and reference electrode with 300 μm gap between them, thus making this a low-cost and disposable device for point-of-care. This sensor has the capability of detecting NA under dynamic pH changes ranging from 9.5 to 6.2 corresponding to before and after NA amplification, respectively. We compared the performance of the developing chip with ISFET electrodes (Microsens, Switzerland) in detection of H^+^ and found correlation coefficient, r = 0.91, between two methods. Based on time response of the sensor and reaction velocity of RPA, the developing sensor can discriminate bacterial strains from culture media in 20 min, indicating capabilities in detecting H^+^ from RPA on disposable chips for pathogen detections. The results show that the micro-planar IrO_2_ was a reliable hydrogen ion sensor and affordability of this design for point-of-care devices.

## 2. Materials and Methods

Iridium tetrachloride of 99.95% purity (Thermo Scientific, USA), 99.99% of purity platinum wire electrode 1.0 mm in diameter (Basinc, USA). Oxalic acid dihydrate, Nafion solution (5 wt%), and 4-(2-hydroxyethyl)-1-piperazineethanesulfonic acid (HEPES) were purchased from Aldrich Sigma. DNA oligonucleotides were synthesized by Integrated DNA Technologies (Irvine, USA) and E. *coli* bacteria were from Real Biotech (Taiwan). TwistAmp® Basic kit was purchased from TwistDX (Cambridge, UK). All other chemicals were of the analytical reagent grade. All solutions were prepared with water from a MilliQ water 7000 system (Millipore, USA). Specific primers were retrieved from the internal transcribed spacer (ITS) sequence of ribosome RNA genes (NCBI: FJ961336) [27] are highly variable and useful for bacteria strain identification [27-29]. Primer for RPA assay were designed based on the consensus sequence according to the guidelines of TwistAmp® DNA amplification kit.

### 2.1. Anodic electrodeposition of IrO_2_ film on gold electrodes

The electrodeposition of IrO_2_ film was performed according to the method described by Yamanaka and our previous studies [12, 14]. The protocol was prepared as follows: 100 mg of iridium tetrachloride was dissolved in 100 mL of water and stirred for 30 min, then 1.0 mL of 30% hydrogen peroxide solution was added and continued to stir for 10 min, and lastly 300 mg of oxalic acid dihydrate was added and continued to stir for 10 min while the final pH 10.5 was adjusted slowly by anhydrous potassium carbonate. The solution was covered and left at room temperature for 2 days for stabilization until resulted dark blue solution was formed. The standard three-electrode system was used: (i) A pair of gold (Au) electrode, including two electrodes, whereas one electrode was used as Ag/AgCl reference electrode, another one was used as the substrate for IrO_2_ deposition, (ii) Platinum (Pt) wire was used as cathode, and (iii) Silver/ Silver chloride (Ag/AgCl) was used as the reference. This approach provides the capability of fabricating a closed pair of planar sensors, one for working electrode (WE) and another for reference electrode. Voltammograms depict the deposition current result from typical coating parameters described in our previous studies [14]. 0.3 × 0.3 mm^2^ electrodes were electrodeposited at 100 mV/s between 0.7 and -0.8 V for 50 to 100 cycles. A blue deposit on planar gold electrodes was observed within 10 min under 85 -120 μA/mm^2^. Thickness growth of IrO_2_ was controlled by increasing current maximums until peaking in the range of 85 -120 μA/mm^2^. Subsequently, pairs of electrodes, one worked as WE and another was coated by Ag/AgCl past for reference electrode, were produced and used for solid-state pH measurements. To complete the construction of the pH electrode, 1.0 µL of 5% Nafion solution was deposited onto the electrode surface and vacuum-dried for 10 min. The electrode pairs were inserted into the chip (**Fig. 1a&b**) and connected to electrical wire using conductive adhesive (Creative materials, USA). The electrical connection was then insulated with silicone rubber (Dow Corning 3140 RTV, USA). The electrodes were calibrated in pH 7.0 solution (Aldrich Sigma, USA) for 2 days to stabilize the potential readings. The pH values are calculated by Nernst equation which is extensively reported in previous papers [9, 10, 14]. In ISFET sensor, pH values are displayed directly through the computer-connected interface electronics to drive the sensor and read the sensor output, provided by Microsens.

**Fig. 1.**
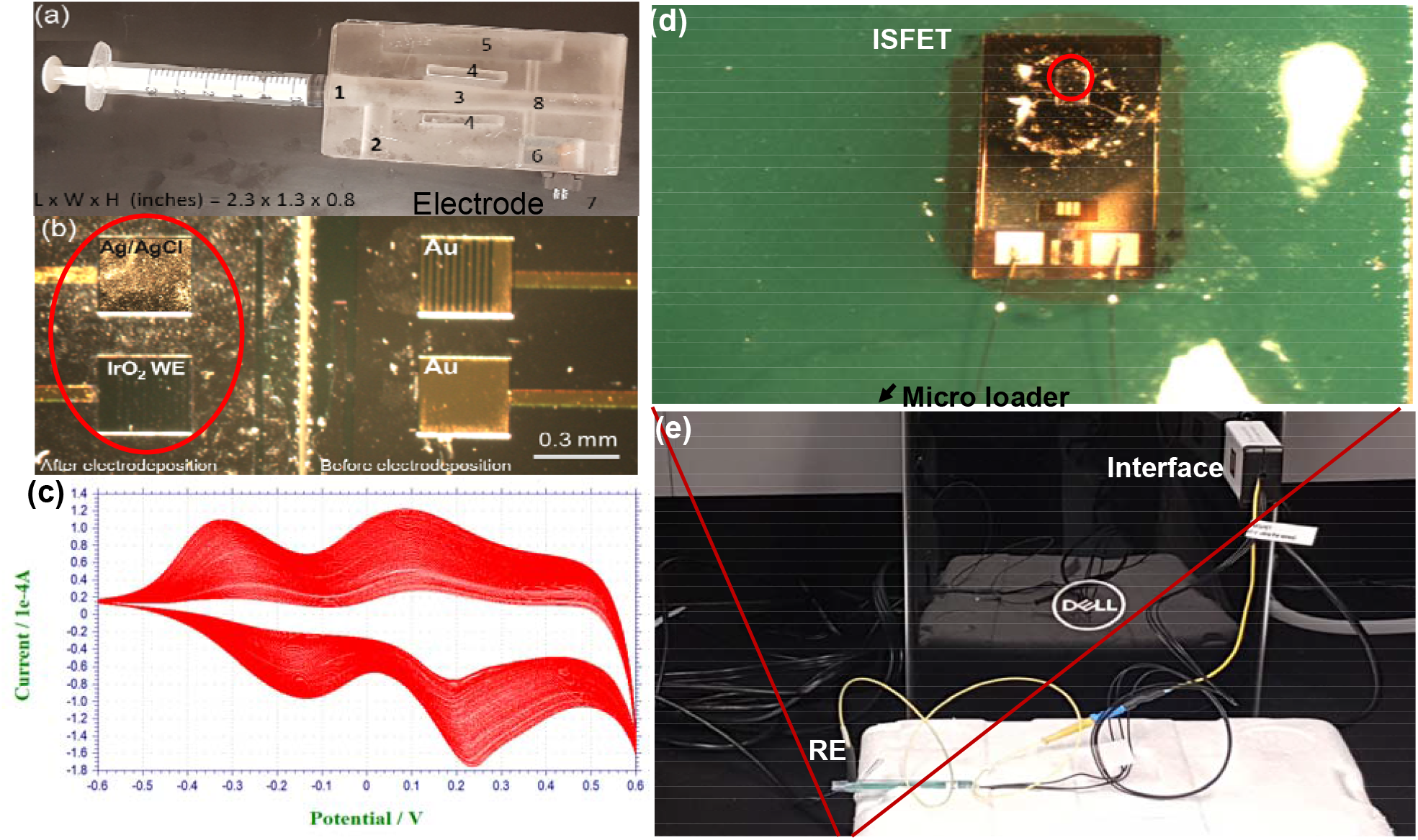
Sensor fabrication for detection of hydrogen ions from nucleic acid amplification. (a) a simple chip for sample loading integrated with IrO2 electrodes. (b) A pair of IrO2 sensing electrode and Ag/AgCl reference electrode, left – before electrodeposition, right – after IrO2 deposition and pseudo-Ag/AgCl reference electrode coated by Ag/AgCl paste. Red circle is a liquid blocker made by Pap Pep that can store 50 µL of samples. (c) cyclic voltammetry spectroscopy of IrO2 electrodeposition. (d) Microscopic image of MSFET 3330 pH-ISFET sensor purchased from Microsens and (e) micro flow cells connected to an interface to measure pH changes.

### 2.2. Electrode Selectivity for ion interferences

The selectivity of the pH electrode was assessed under DNA amplification conditions. Before measurement, 50 µL of pH 7.0 solution was used for potential stabilization. Subsequently, 50 µL of each PCR buffers containing different ions was dropped on the electrode. The changes in millivolt reading were recorded. The interference effects of Mg^2+^, SO_4_ ^2+^, and NH^4+^ in PCR and RPA buffers were evaluated. The effect of dissolved oxygen was eliminated by leaving the samples in vacuum chamber before measurement.

### 2.3. pH calibration and released hydrogen ions measurements

The electrodes were calibrated for their pH response in a series of a calibrated standard pH buffer (pH 4, 7, and 10). Calibration curves cover the PCR and RPA buffers. The PCR and RPA were both evaluated for their pH responses in a series of DNA inputs in the pH range 6.2 – 7.0. The pH of DNA amplification buffers before reactions were identified at 8.5 – 9.2. Through potential measurement using IrO_2_ electrode, the number of released H^+^ from DNA amplifications were proportional to the number of DNA copies or amplification cycles, which indicates pH values are shifted to the acid side depending on the concentration of target. The reproducibility of the pH responses in the pH range was 6.2 – 9.2 evaluated by changing PCR cycles and time of DNA amplifications in RPA. The conventional detection using SYBR Green staining were conducted to observe increased DNA concentration after DNA amplifications.

Data received from PCR and RPA were down sampled to one data point for 5 cycles and one data point per minute, respectively. Sensor drift due to the low DNA copy number at some first cycles of the DNA amplifications was corrected by curve fitting the data from each electrical signal using data points from the first 5 cycles or 3 minutes of PCR and RPA, respectively. To remove ISFET background signals, ISFET signals (ΔmV) calculated from the reference chamber were subtracted from ISFET signals derived from the experimental chambers. The curves were normalized by dividing each ΔmV value by the maximum ΔmV value and plotted against time or cycle.

### 2.4. Validation the developing sensor by application of the ISFET method

The ISFET sensor (MSFET 3330 pH-ISFET sensor) used in this study was purchased from Microsens, Switzerland and consisted of three terminals, source, drain and a reference electrode substituting the metal gate. Tantalum pentoxide (Ta_2_O_5_) made up the gate insulator and the reference electrode were of miniature Ag/AgCl reference electrode (MSREF). A MSFET interface, provided by Microsens, was used to drive the sensor and read the sensor output. The ISFET was placed into the bottom of custom-made flow cell with the reference electrode adapted in the top of the compartment that had a volume reaction of 50 µL. 50 µL of DNA amplification samples was applied to the flow cells through a micro loader (Eppendorf, USA). The output voltage was proportional to the pH of the DNA samples with a typical sensitivity of 55 mV/pH. Performance of the device was routinely assessed using standard buffers of pH values 4, 7, and 10 (Aldrich Sigma, USA).

## 3. Results and discussion

### 3.1. Anodic deposition of Iridium oxide and device setup

Cyclic voltammograms for the IrO_2_ electrodeposition on bare gold electrodes show reversible redox behavior of the activated iridium electrodes. **Fig. 1c** shows the response of three fabricated sensors (average values) with a surface area of 0.3 ×0.3 mm^2^, produced with 100 sweeps at 100 mV/s, exhibiting a super-Nernstian average of 59.6 mV/pH and ranging from 55.3 to 67.8 mV/pH (confidence interval (CI) 96%, n= 7 samples). In previous studies, the generated potential between the working and reference electrodes strongly affects the kinetics of reactions occurring on the IrO_2_ surface is reported [30]. Yamanaka reported that the mechanism by which IrO_2_ was formed involves anodic oxidation of the oxalate ligand to CO_2_ with concomitant deposition of iridium as iridium (IV) oxide at the anode [12]. The fabricating sensor showed the characteristic electrochromic behavior of IrO_2_ with alkaline iridium (IV) solution containing oxalate, which prevents iridium precipitation in the alkaline medium by complex formation during redox reaction. The redox reaction occurred on the WE through the ion-exchange mechanism. The WE were made from depositing IrO_2_ on the top of an Au substrate, the most promising proton exchange membrane electrolyzer electrocatalyst [31], is formed by H^+^ and/or OH^−^ ions from solution, accumulating at the interface of IrO_2_ electrode (**Fig. 1b**). After each PCR cycle, changes in the number of released hydrogen ions, then, lead to changes in pH value, which makes ratio H^+^/OH^-^ in the surface layer of IrO_2_ electrode. The ISFET sensor (MSFET 3330 pH-ISFET sensor) (**Fig.1d**) connected with the interface was shown in **Fig. 1e**.

### 3.2. Detection of DNA amplifications on iridium oxide electrode

The detection principle was described in **Fig. 2 (a-c)** including RPA reaction (**Fig. 2a**), hydrogen ion detection (**Fig. 2b**) by pH sensor (upper) through open circuit potential (OCP) and ISFET (lower) and converting the chemical signal (H+) to electrical signals (**Fig. 2c**). In pH sensor, IrOx electrodes detect the pH change due to accumulating released H^+^ from DNA amplifications and converts the chemical signals (H+) into electrical signals. The DNA amplification is initiated by binding of specific primers (**Table 1**) and successive incorporation of nucleotides leads to the liberation of H^+^ and consequently a decrease in pH of the reaction [2]. With mismatch primers at 3′-OH end or unspecific primers, the amplification does not proceed to produce hydrogen ions, which do not result in a significant pH change.

**Table 1.**
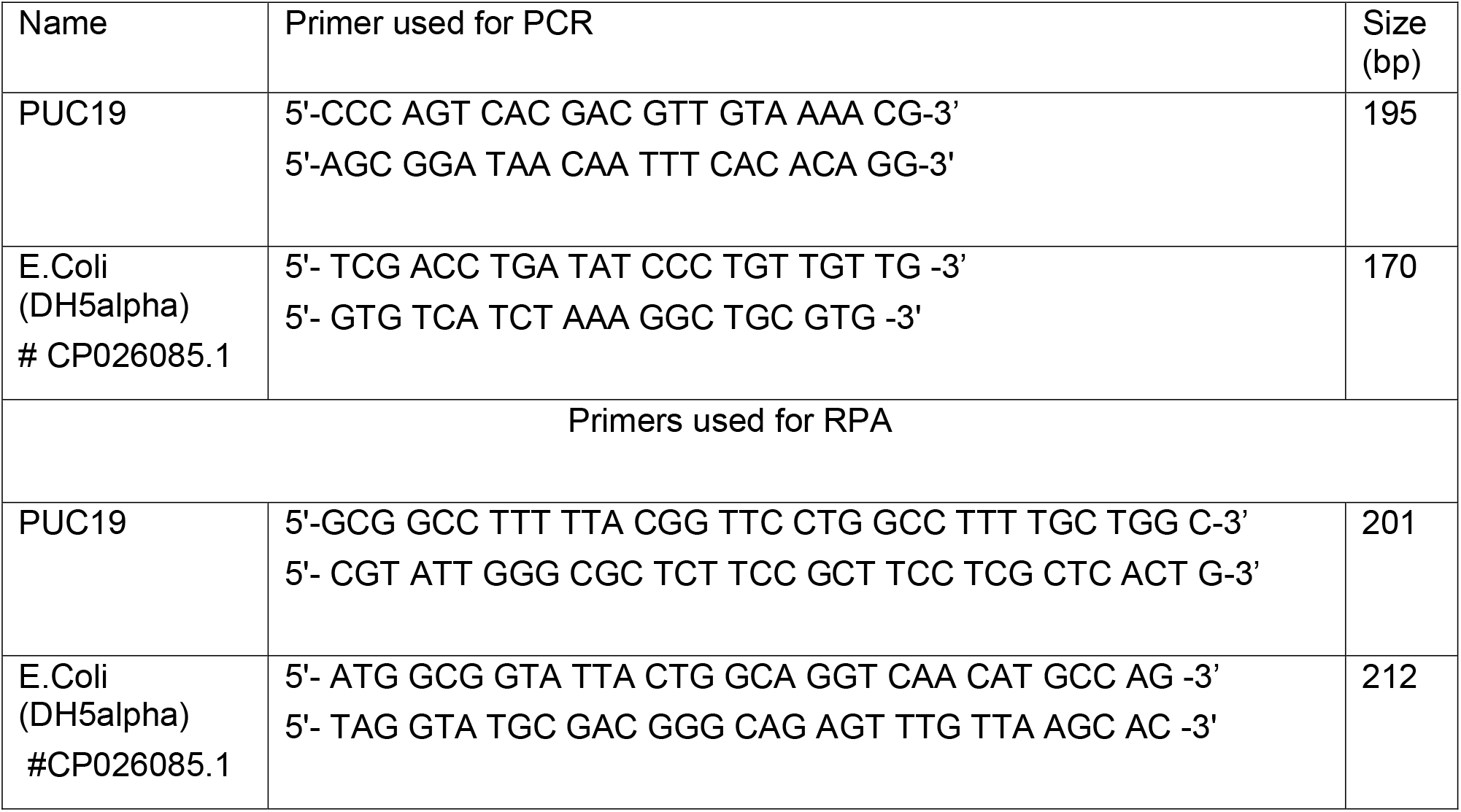
Primer sequences used for PCR and RPA.

**Fig. 2.**
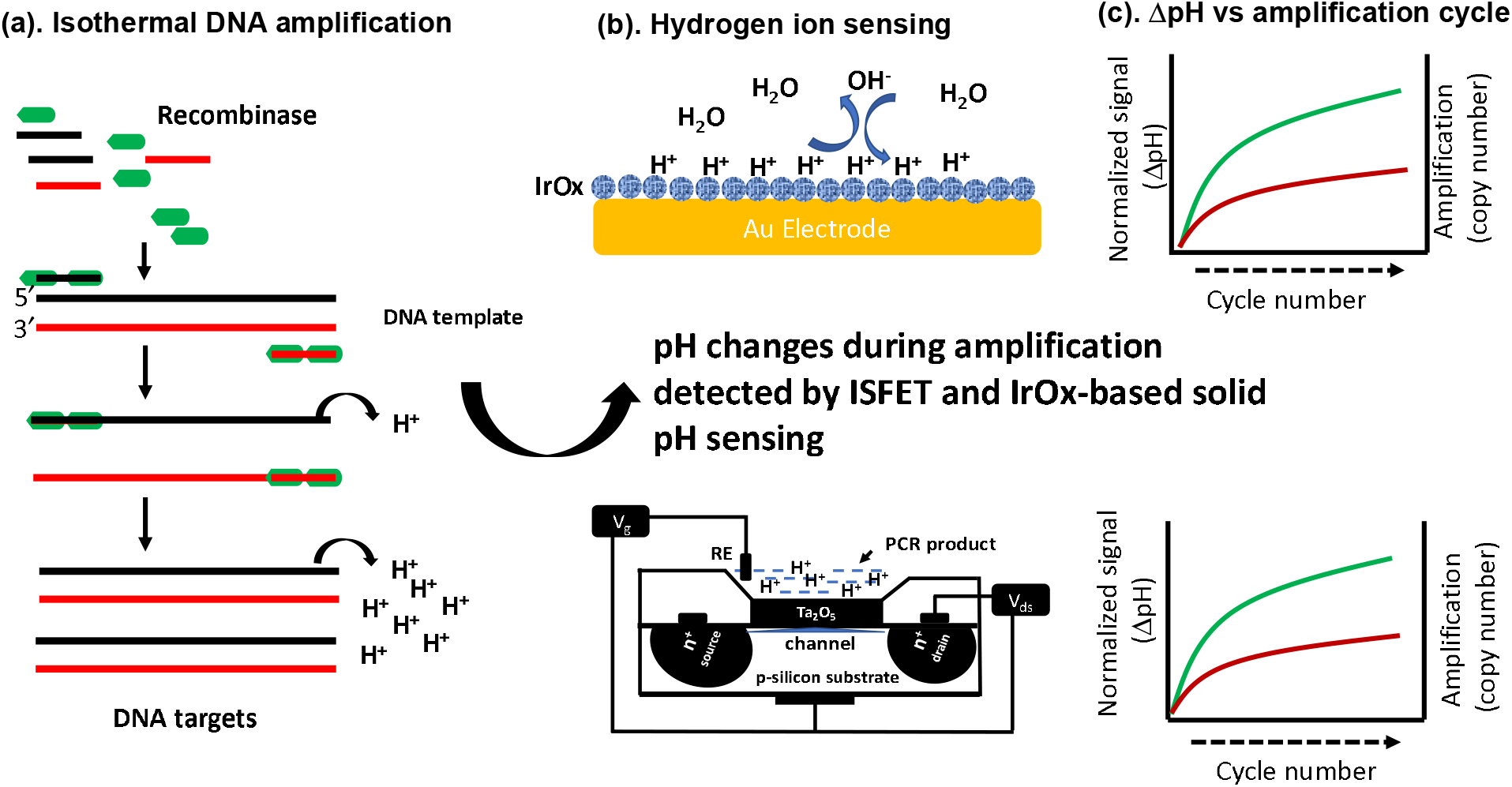
Principle of schematic detection. (a) Hydrogen ions are released during the RPA process. When one nucleotide molecule is integrated into extending DNA backbones, one hydrogen ion is released into the solution. (b) Released hydrogen ions that lead to pH change are detected by IrO2 (upper) and ISFET sensor (lower). (c) Schematic detection of electrical signal converted from chemical signal.

We optimized the DNA amplification conditions for two well-known amplification techniques, PCR and RPA, under low-ion conditions while keeping amplification efficiency and specificity [5]. The releases H^+^ has been used as a reliable indicator for DNA amplification. Using PUC19 as a reference target, we run DNA amplifications with specific primers (**Table 1**) in a standard thermal cycler (MaxyGene II, Corning Axygen). pH analysis for amplified DNA samples were measured after each 5 cycles using the developing IrO_2_ electrode (**Fig. 3a, red line**) and the ISFET-based pH probe (**Fig. 3a, black line**. In RPA, released H^+^ was detected through a time course of 15 min was shown in **Fig. 3b**. The IrO_2_ sensor (**Fig. 3b**, red line) showed a relevant normalized ΔpH values to that of ISFET sensor (**Fig. 3b**, black line). The average difference (0.12 unit) recorded between IrO_2_ sensors and ISFET sensors was could be due to high presence of Mg^2+^ in RPA buffer [32]. The quantity of amplified DNA was measured by qPCR and conventional PCR (**Fig. 3c**). A linear association (R^2^ ∼ 0.978) between the PCR-based amplification yield and pH change was observed (**Fig. 3d, red line**). Similarly, RPA reaction shows a strong correlation (R^2^ = 0.956) between the pH values and the number of DNA during the reaction in 15 min (**Fig. 3d, black line**). Specific DNA amplification by agarose gel electrophoresis with a single PCR product were observed in insets of **Fig. 3a-c**.

**Fig. 3.**
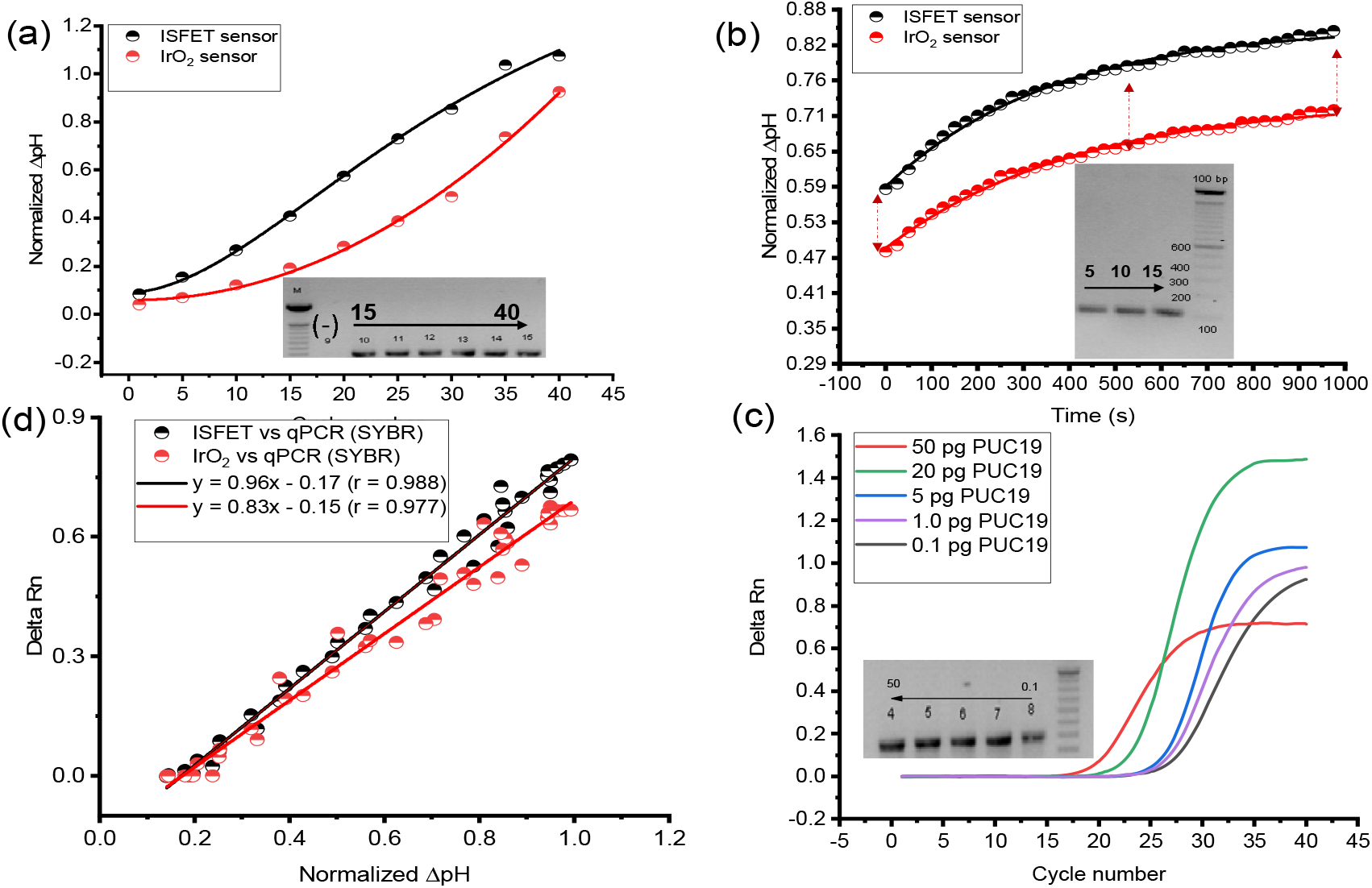
Performance of the sensing platform. pH changes of nucleic acid samples amplified from PCR (a) and RPA (b) and detected by IrO2 (red line) and ISFET sensor (black line). (c) qPCR results of different DNA concentrations of PUC19. (d) Correlation between sensor platforms and qPCR method; IrO2 sensor versus qPCR (red line) and ISFET sensor versus qPCR (black line).

Performance of the sensing platform. pH changes of nucleic acid samples amplified from PCR (a) and RPA (b) and detected by IrO2 (red line) and ISFET sensor (black line). (c) qPCR results of different DNA concentrations of PUC19. (d) Correlation between sensor platforms and qPCR method; IrO2 sensor versus qPCR (red line) and ISFET sensor versus qPCR (black line).

The pH change from PCR-based DNA amplifications from PUC19 was demonstrated by the progressive increase of differential potential from cycle 25 (**Fig. 4a**) and the threshold value of sensing platforms, relevant to qPCR, showed the correlation between number of copies to threshold values (**Fig. 4b**). The sensitivity and the potential quantitative application of IrO_2_, a solid-state pH sensor, were determined through ITS gene of E. coli (NCBI: FJ961336) A tenfold-serial dilution of *E. coli* genome was used to compare the performance of the sensing method with conventional SYBR gene-based qPCR (Thermo Scientific, USA). All the IrO_2_, ISFET sensing, and the optical platforms resulted in a comparable efficiency of PCR, 90.2%, 94.8%, and 97.3%, respectively. Good coefficient correlations were achieved at 0.906, 0.933, and 0.998 for iridium oxide, ISFET, and qPCR, respectively. Lowest dilution of the number of DNA was detected in all platforms were 25 copies of genomic DNA (**Fig. 4b**).

**Fig. 4.**
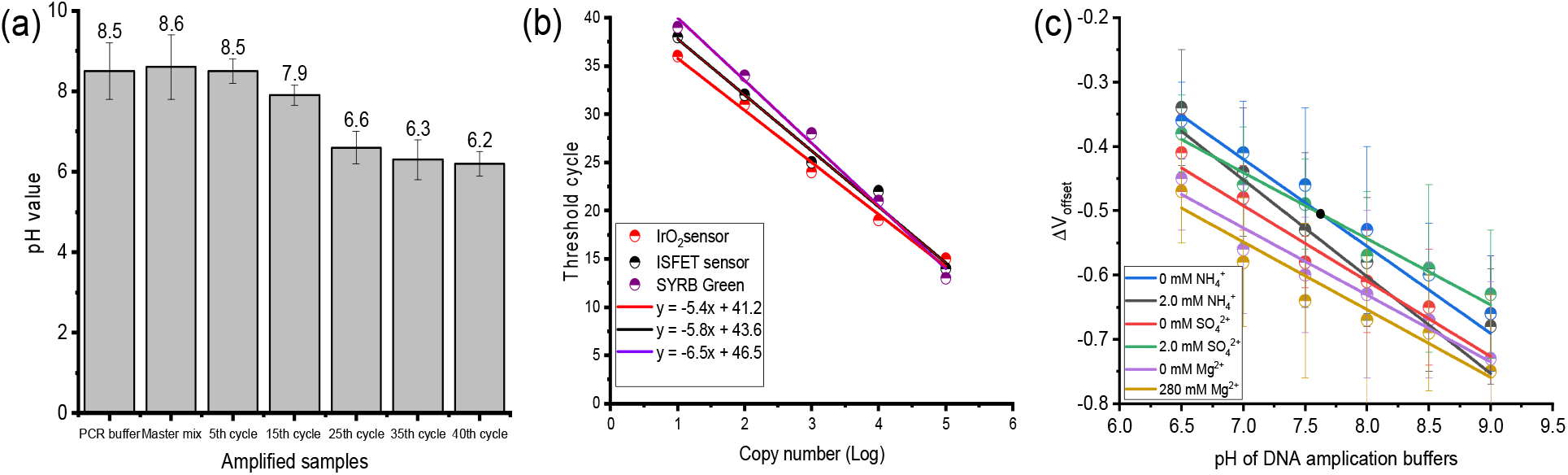
pH changes during DNA amplification cycles and ion interferences. (a) pH value changes from bare PCR buffers to amplified samples after 40 cycles. pH values reduced from 8.5 to 6.2 are observed. (b). The correlation between DNA copies and threshold values. DNA copies were identified by Nanodrop method. (c) Ion interferences were observed in the IrO2 sensor with an average difference of 0.05.

In addition, ion interferences of Mg^2+^, NH_4_^+^, and SO_4_^2+^ presenting in DNA amplification buffers were shown in **Fig. 4c**. The variability of pH change due to ion interference was calculated at average 0.05 pH. The previous studies reported that IrO_2_ electrodes showed very low sensitivity for difference ion species [33]. However, the sensor surface may be further studied to attenuate the interference of anionic redox species, especially for Mg^2+^. Together these data indicate that IrO_2_ sensing can be a sensitivity and reliability for DNA amplification.

### 3.3. pH-sensitive amplification by RPA for *E. coli* bacteria

Amplification and detection of *E. coli* bacteria based on specific primers were shown in **Table 1**. We used super Pap Pep, a liquid blocker, to draw a circle around a pair of electrodes to make 50-µl chamber flow cell. The chip was connected to a Palmsens 4 device (Palmsens, Netherland) to do potential measurement.

To experimentally establish the real-time performance of isothermal DNA amplification on the solid-state pH sensor, we measured released hydrogen ions from RPA reactions. We dispensed RPA amplification reagents containing primers targeting *E. coli* strains including DH5alpha, 9 BL21, 3 HB101, 6 TOP10, and 3 JM109 and purified genomic DNA into hydrophobic circle on a chamber while another chamber was used as a reference control reaction with no template. 50 µL of a final RPA assay was run on the chip with 3 µL of genomic DNA, according to the operating manual of TwistAmp Basic and other previous papers [21, 24, 26]. A shaking incubator was used to perform the reaction at 39 °C for 10 min with 100 rpm. The change of pH signal was monitored and shown in **Fig. 5a**, referenced by PCR method in **Fig. 5b**. The amplification curves of the IrO_2_ sensing measured by change in potential were highly comparable to the fluorescence signal detected by the qPCR [2]. In addition, a fraction of each reaction mixture on an agarose gel and observed only one specific band in the amplification reactions, whereas no bands were observed in the no-template control reactions. The results of gel electrophoresis, slope of thermal cycler, and hydrogen ions sensing platform were comparable and reliable, suggesting that IrO_2_ platform have similar performance with ISFET and qPCR techniques in detection of DNA amplification.

**Fig. 5.**
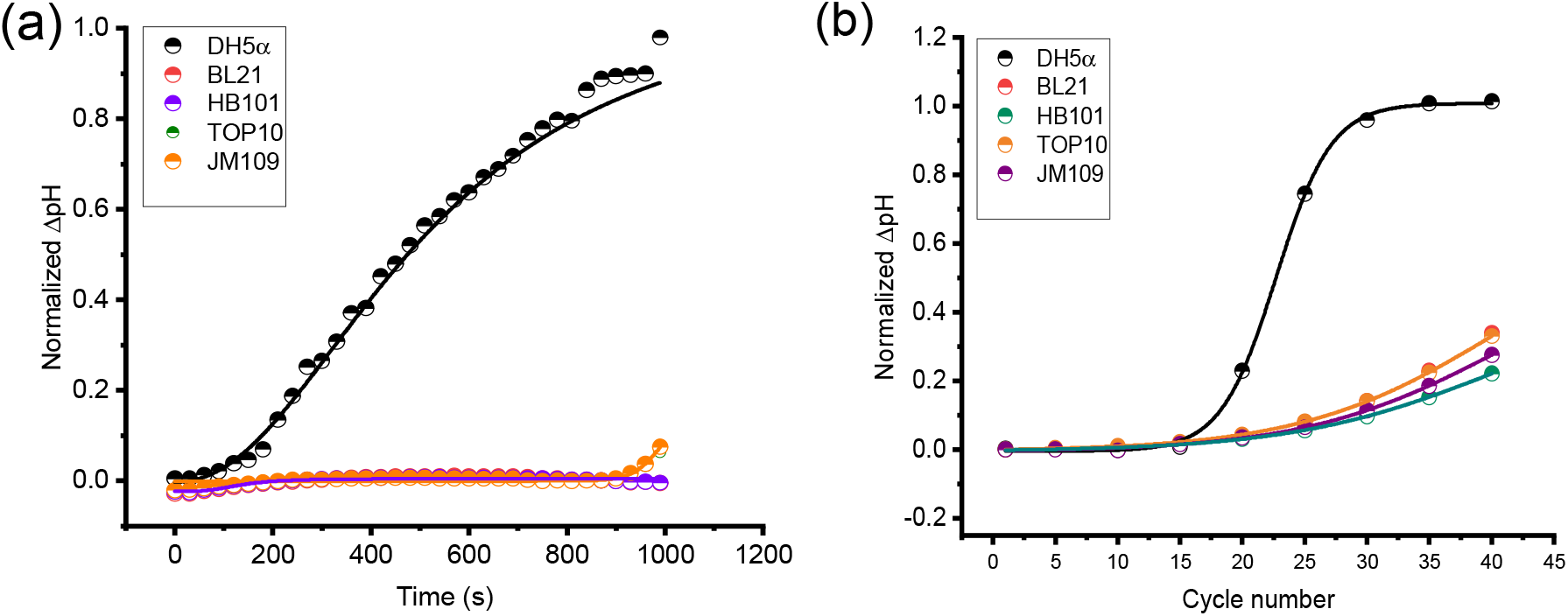
Specificity of the sensor in detection of E. coli strains. The specificity of the sensors in this experiment is highly dependent on primer specificity used in DNA amplification. The specific primers for DH5alpha were used. The sensor detected DH5alpha from other E. coli strains by RPA (a) and PCR (b).

## 4. Conclusions

Overall, the amplification curves between SYBR-based qPCR and electrical signal of the developing chip are comparable. Approximate 25 genomic copies were detected with PCR and RPA reaction on the pH sensing platform. The versatility of the electrical platform and the capacity to quantitate provides an opportunity for creating multiple chips at high-density to multiple pathogen detection. However, the slight variability of pH changes observed by batch-to-batch IrO_2_ film. The variability may be from chemical surface of iridium oxide electrode. Such chemical variability can be minimized by chemical surface modification in future studies. The advantage of the electrical sensing platform is lowest cost, less contamination, same sensitivity with optical platforms, and upgrading to multiple detection with lowest cost by using paper-based electrodes, which is advantageous in the development of a portable and multiple platforms for pathogen detection based on nucleic acid amplification. The simple open circuit can be miniaturized to integrate in the point-of-care apparatus. For example, we will increase the number of sensing electrodes and integrate the sensing chamber into a genome isolation module to develop a full device.

## Author Contributions

Conceptualization

Anh H. Nguyen, Hung Cao

Data curation

Samir Malhotra, Anh H. Nguyen

Formal analysis

Samir Malhotra, Anh H. Nguyen,

Investigation

Samir Malhotra, Anh H. Nguyen,

Methodology

Samir Malhotra, Anh H. Nguyen, Hung Cao

Writing – original draft

Samir Malhotra, Anh H. Nguyen.

Writing – review & editing

Samir Malhotra, Anh H. Nguyen, Hung Cao

## Acknowledgement

The authors would like to acknowledge the financial support from the NSF CAREER Award #1917105 (H.C.), the NSF #1936519 (J.L. and H.C), the NIH SBIR grant #R44OD024874 (M.P.H.L and H.C.), the NIH HL107304 and HL081753 (X.X.).

## References

1. Leonardo, S.; Toldrà, A.; Campàs, M., Biosensors Based on Isothermal DNA Amplification for Bacterial Detection in Food Safety and Environmental Monitoring. Sensors (Basel) 2021, 21, (2).

2. Toumazou, C.; Shepherd, L. M.; Reed, S. C.; Chen, G. I.; Patel, A.; Garner, D. M.; Wang, C. J.; Ou, C. P.; Amin-Desai, K.; Athanasiou, P.; Bai, H.; Brizido, I. M.; Caldwell, B.; Coomber-Alford, D.; Georgiou, P.; Jordan, K. S.; Joyce, J. C.; La Mura, M.; Morley, D.; Sathyavruthan, S.; Temelso, S.; Thomas, R. E.; Zhang, L., Simultaneous DNA amplification and detection using a pH-sensing semiconductor system. Nat Methods 2013, 10, (7), 641–6.

3. Hua, X.; Yang, E.; Yang, W.; Yuan, R.; Xu, W., LAMP-generated H(+) ions-induced dimer i-motif as signal transducer for ultrasensitive electrochemical detection of DNA. Chem Commun (Camb) 2019, 55, (83), 12463–12466.

4. Park, S. H.; Aydin, M.; Fan, P.; Lee, S.; Teng, L.; Kim, S. A.; Ahn, S.; Ricke, S. C.; Shi, Z.; Jeong, K. C., Chapter 16 - Detection Strategies for Foodborne Salmonella and Prospects for Utilization of Whole Genome Sequencing Approaches. In Food and Feed Safety Systems and Analysis, Ricke, S. C.; Atungulu, G. G.; Rainwater, C. E.; Park, S. H., Eds. Academic Press: 2018; pp 289–308.

5. Lee, K.-H.; Lee, D.; Yoon, J.; Kwon, O.; Lee, J., A Sensitive Potentiometric Sensor for Isothermal Amplification-Coupled Detection of Nucleic Acids. Sensors 2018, 18, (7), 2277.

6. Pourmand, N.; Karhanek, M.; Persson, H. H. J.; Webb, C. D.; Lee, T. H.; Zahradníková, A.; Davis, R. W., Direct electrical detection of DNA synthesis. Proceedings of the National Academy of Sciences 2006, 103, (17), 6466–6470.

7. Rothberg, J. M.; Hinz, W.; Rearick, T. M.; Schultz, J.; Mileski, W.; Davey, M.; Leamon, J. H.; Johnson, K.; Milgrew, M. J.; Edwards, M.; Hoon, J.; Simons, J. F.; Marran, D.; Myers, J. W.; Davidson, J. F.; Branting, A.; Nobile, J. R.; Puc, B. P.; Light, D.; Clark, T. A.; Huber, M.; Branciforte, J. T.; Stoner, I. B.; Cawley, S. E.; Lyons, M.; Fu, Y.; Homer, N.; Sedova, M.; Miao, X.; Reed, B.; Sabina, J.; Feierstein, E.; Schorn, M.; Alanjary, M.; Dimalanta, E.; Dressman, D.; Kasinskas, R.; Sokolsky, T.; Fidanza, J. A.; Namsaraev, E.; McKernan, K. J.; Williams, A.; Roth, G. T.; Bustillo, J., An integrated semiconductor device enabling non-optical genome sequencing. Nature 2011, 475, (7356), 348–352.

8. Chaisiwamongkhol, K.; Batchelor-McAuley, C.; Compton, R. G., Optimising amperometric pH sensing in blood samples: an iridium oxide electrode for blood pH sensing. Analyst 2019, 144, (4), 1386–1393.

9. Zhou, L.; Cheng, C.; Li, X.; Ding, J.; Liu, Q.; Su, B., Nanochannel Templated Iridium Oxide Nanostructures for Wide-Range pH Sensing from Solutions to Human Skin Surface. Analytical Chemistry 2020, 92, (5), 3844–3851.

10. Prats-Alfonso, E.; Abad, L.; Casañ-Pastor, N.; Gonzalo-Ruiz, J.; Baldrich, E., Iridium oxide pH sensor for biomedical applications. Case urea-urease in real urine samples. Biosens Bioelectron 2013, 39, (1), 163–9.

11. Tabata, M.; Katayama, Y.; Mannan, F.; Seichi, A.; Suzuki, K.; Goda, T.; Matsumoto, A.; Miyahara, Y., Label-free and Electrochemical Detection of Nucleic Acids Based on Isothermal Amplification in Combination with Solid-state pH Sensor. Procedia Engineering 2016, 168, 419–422.

12. Yamanaka, K., Anodically Electrodeposited Iridium Oxide Films (AEIROF) from Alkaline Solutions for Electrochromic Display Devices. Japanese Journal of Applied Physics 1989, 28, (Part 1, No. 4), 632–637.

13. Ng, S. R.; O’Hare, D., An iridium oxide microelectrode for monitoring acute local pH changes of endothelial cells. Analyst 2015, 140, (12), 4224–4231.

14. Marsh, P.; Manjakkal, L.; Yang, X.; Huerta, M.; Le, T.; Thiel, L.; Chiao, J. C.; Cao, H.; Dahiya, R., Flexible Iridium Oxide Based pH Sensor Integrated With Inductively Coupled Wireless Transmission System for Wearable Applications. IEEE Sensors Journal 2020, 20, (10), 5130–5138.

15. Tarlov, M. J.; Semancik, S.; Kreider, K. G., Mechanistic and response studies of iridium oxide pH sensors. Sensors and Actuators B: Chemical 1990, 1, (1), 293–297.

16. Kurzweil, P., Metal Oxides and Ion-Exchanging Surfaces as pH Sensors in Liquids: State-of-the-Art and Outlook. Sensors (Basel) 2009, 9, (6), 4955–85.

17. Zea, M.; Moya, A.; Fritsch, M.; Ramon, E.; Villa, R.; Gabriel, G., Enhanced Performance Stability of Iridium Oxide-Based pH Sensors Fabricated on Rough Inkjet-Printed Platinum. ACS Applied Materials & Interfaces 2019, 11, (16), 15160–15169.

18. Xi, Y.; Guo, Z.; Wang, L.; Xu, Q.; Ruan, T.; Liu, J., Fabrication and Characterization of Iridium Oxide pH Microelectrodes Based on Sputter Deposition Method. Sensors (Basel, Switzerland) 2021, 21, (15), 4996.

19. Cialone, M.; Fernandez-Barcia, M.; Celegato, F.; Coisson, M.; Barrera, G.; Uhlemann, M.; Gebert, A.; Sort, J.; Pellicer, E.; Rizzi, P.; Tiberto, P., A comparative study of the influence of the deposition technique (electrodeposition versus sputtering) on the properties of nanostructured Fe(70)Pd(30) films. Sci Technol Adv Mater 2020, 21, (1), 424–434.

20. Daher, R. K.; Stewart, G.; Boissinot, M.; Bergeron, M. G., Recombinase Polymerase Amplification for Diagnostic Applications. Clinical chemistry 2016, 62, (7), 947–58.

21. Archer, J.; Barksby, R.; Pennance, T.; Rostron, P.; Bakar, F.; Knopp, S.; Allan, F.; Kabole, F.; Ali, S. M.; Ame, S. M.; Rollinson, D.; Webster, B. L., Analytical and Clinical Assessment of a Portable, Isothermal Recombinase Polymerase Amplification (RPA) Assay for the Molecular Diagnosis of Urogenital Schistosomiasis. Molecules 2020, 25, (18).

22. Zhao, G.; Hou, P.; Huan, Y.; He, C.; Wang, H.; He, H., Development of a recombinase polymerase amplification combined with a lateral flow dipstick assay for rapid detection of the Mycoplasma bovis. BMC Vet Res 2018, 14, (1), 412–412.

23. Babujee, L.; Witherell, R. A.; Mikami, K.; Aiuchi, D.; Charkowski, A. O.; Rakotondrafara, M., Optimization of an isothermal recombinase polymerase amplification method for real-time detection of Potato virus Y O and N types in potato. J Virol Methods 2019, 267, 16–21.

24. Ma, Q.; Yao, J.; Yuan, S.; Liu, H.; Wei, N.; Zhang, J.; Shan, W., Development of a lateral flow recombinase polymerase amplification assay for rapid and visual detection of Cryptococcus neoformans/C. gattii in cerebral spinal fluid. BMC Infect Dis 2019, 19, (1), 108.

25. Geng, Y.; Liu, G.; Liu, L.; Deng, Q.; Zhao, L.; Sun, X. X.; Wang, J.; Zhao, B.; Wang, J., Real-time recombinase polymerase amplification assay for the rapid and sensitive detection of Campylobacter jejuni in food samples. J Microbiol Methods 2019, 157, 31–36.

26. Zaghloul, H.; El-Shahat, M., Recombinase polymerase amplification as a promising tool in hepatitis C virus diagnosis. World J Hepatol 2014, 6, (12), 916–922.

27. Magray, M. S.; Kumar, A.; Rawat, A. K.; Srivastava, S., Identification of Escherichia coli through analysis of 16S rRNA and 16S-23S rRNA internal transcribed spacer region sequences. Bioinformation 2011, 6, (10), 370–1.

28. da Cunha, M. d. L. R. d. S., Molecular Biology in Microbiological AnalysislZI. In Reference Module in Food Science, Elsevier: 2019.

29. Man, S. M.; Kaakoush, N. O.; Octavia, S.; Mitchell, H., The internal transcribed spacer region, a new tool for use in species differentiation and delineation of systematic relationships within the Campylobacter genus. Appl Environ Microbiol 2010, 76, (10), 3071–3081.

30. Kuo, D.-Y.; Kawasaki, J. K.; Nelson, J. N.; Kloppenburg, J.; Hautier, G.; Shen, K. M.; Schlom, D. G.; Suntivich, J., Influence of Surface Adsorption on the Oxygen Evolution Reaction on IrO2(110). Journal of the American Chemical Society 2017, 139, (9), 3473–3479.

31. Jovanovič, P.; Hodnik, N.; Ruiz-Zepeda, F.; Arčon, I.; Jozinović, B.; Zorko, M.; Bele, M.; Šala, M.; Šelih, V. S.; Hočevar, S.; Gaberšček, M., Electrochemical Dissolution of Iridium and Iridium Oxide Particles in Acidic Media: Transmission Electron Microscopy, Electrochemical Flow Cell Coupled to Inductively Coupled Plasma Mass Spectrometry, and X-ray Absorption Spectroscopy Study. Journal of the American Chemical Society 2017, 139, (36), 12837–12846.

32. Wang, S.; McDonnell, E. H.; Sedor, F. A.; Toffaletti, J. G., pH effects on measurements of ionized calcium and ionized magnesium in blood. Archives of pathology & laboratory medicine 2002, 126, (8), 947–50.

33. Marzouk, S. A.; Ufer, S.; Buck, R. P.; Johnson, T. A.; Dunlap, L. A.; Cascio, W. E., Electrodeposited iridium oxide pH electrode for measurement of extracellular myocardial acidosis during acute ischemia. Anal Chem 1998, 70, (23), 5054–61.

